# Structural basis of dynamic P5CS filaments

**DOI:** 10.1101/2021.12.02.470899

**Authors:** Jiale Zhong, Chen-Jun Guo, Xian Zhou, Chia-Chun Chang, Boqi Yin, Tianyi Zhang, Huan-Huan Hu, Guang-Ming Lu, Ji-Long Liu

**Affiliations:** School of Life Science and Technology, ShanghaiTech University, Shanghai, 201210, China; Institute of Biochemistry and Cell Biology, Shanghai Institutes for Biological Sciences, Chinese Academy of Sciences, Shanghai, 200031, China; University of Chinese Academy of Sciences, Beijing, 100049, China

**Keywords:** P5CS, cytoophidium, cryo-EM, filamentation, proline biosynthesis

## Abstract

The bifunctional enzyme Δ^1^-pyrroline-5-carboxylate synthase (P5CS) is central to the synthesis of proline and ornithine. Pathogenic mutations in P5CS gene (ALDH18A1) lead to neurocutaneous syndrome and skin relaxation connective tissue disease in humans, and P5CS deficiency seriously damages the ability to resist adversity in plants, which has an essential role in agriculture and human health. Recently, P5CS has been demonstrated forming the cytoophidium in vivo and filaments in vitro. However, the underlying mechanism for the function of P5CS filamentation and catalyze the synthesis of P5C is hardly accessible without structural basis. Here, we have succeeded in determining the full-length structures of *Drosophila* P5CS filament in three states at resolution from 3.1 to 4.3 Å under cryo-electron microscopy, we observed the distinct ligand-binding states and conformational changes for GK and GPR domain separately. These structures show the distinctive spiral filament is assembled by P5CS tetramers and stabilized by multiple interfaces. Point mutations that deplete such interactions disturb P5CS filamentation and greatly reduce the activity. Our findings reveal a previously undescribed mechanism that filamentation is crucial for the coordination between GK and GPR domains, and provide insights into structural basis for catalysis function of P5CS filament.

## Introduction

Pyrroline-5-carboxylate synthase (P5CS) is crucial for proline and ornithine metabolism(*1–4*). In plants, proline synthesis is associated to plant stress resistance(*4*). In humans, over 30 mutations of P5CS have been identified as the cause of rare diseases(*2–6*). In addition, the glutamine-proline regulatory axis has been regarded as a promising target for cancer treatments(*7, 8*). Therefore, P5CS is of great significance in agriculture and human health.

Previous studies have revealed a distinctive compartmentation of enzymes via filamentation(*9–13*). This filamentous structure is membraneless and termed the cytoophidium for its appearance(*9, 14*). The cytoophidium has emerged as a mechanism for the regulation of several metabolic enzymes(*14, 15*). Recently, we have identified P5CS as a novel cytoophidium-forming enzyme in *Drosophila* tissues and also demonstrated the filamentation of P5CS *in vitro*(*16*).

P5CS corresponds to two individual proteins in prokaryotes and some lower eukaryotes such as yeast. One is the glutamate kinase (GK, *pro*B gene), and the other is gamma-glutamyl phosphate reductase (GPR, *pro*A gene) in *E. coli*. The kinetic analysis suggests that bacterial GK and GPR could form a complex(*17*). The dual functions of P5CS in higher eukaryotes implicate that the GK and GPR have evolved into one single protein for coupling reactions of two domains. However, the full-length structure of P5CS has not yet been reported. The underlying molecular mechanisms of the catalytic reaction and the function of filamentation remain largely unknown.

In this research, by using cryo-electron microscopy, we resolve the full-length structure of *Drosophila melanogaster* P5CS filament in multiple states. Our reconstruction structures at 3.1 to 4.3 Å resolutions provide detailed information of the P5CS filament in different ligand binding conformations. According to the structural basis, we reveal the working mechanisms for the spiral structure of P5CS filaments.

## Results and Discussion

### Overall P5CS filament structures and characterizations

Bifunctional P5CS, which is comprised by two domains, catalyzes the first and second step in the biosynthesis of proline from glutamate. The GK domain catalyzes glutamate phosphorylation and the GPR domain catalyzes the NADPH-dependent reduction of γ-glutamyl phosphate (G5P) to glutamate-γ-semialdehyde (GSA). The end product P5C is formed by a spontaneous cyclisation reaction of GSA (Figure 1A), and will be used in the production of proline. In order to resolve the filament structure of *Drosophila* P5CS, we expressed and purified *Drosophila melanogaster* full-length P5CS proteins, specific details were described in the method. We first analyzed the filamentation of P5CS at its APO state and state with all substrates by negative staining (Figure S1A-C). The results show that P5CS proteins self-assemble into filament without the requirement of ligands, and the addition of substrates could enhance the length of filaments. In consistent with our previous study, L-glutamate, which is a substrate of P5CS, is critical in promoting the formation and stability P5CS filament(*16*).

**Figure 1.**
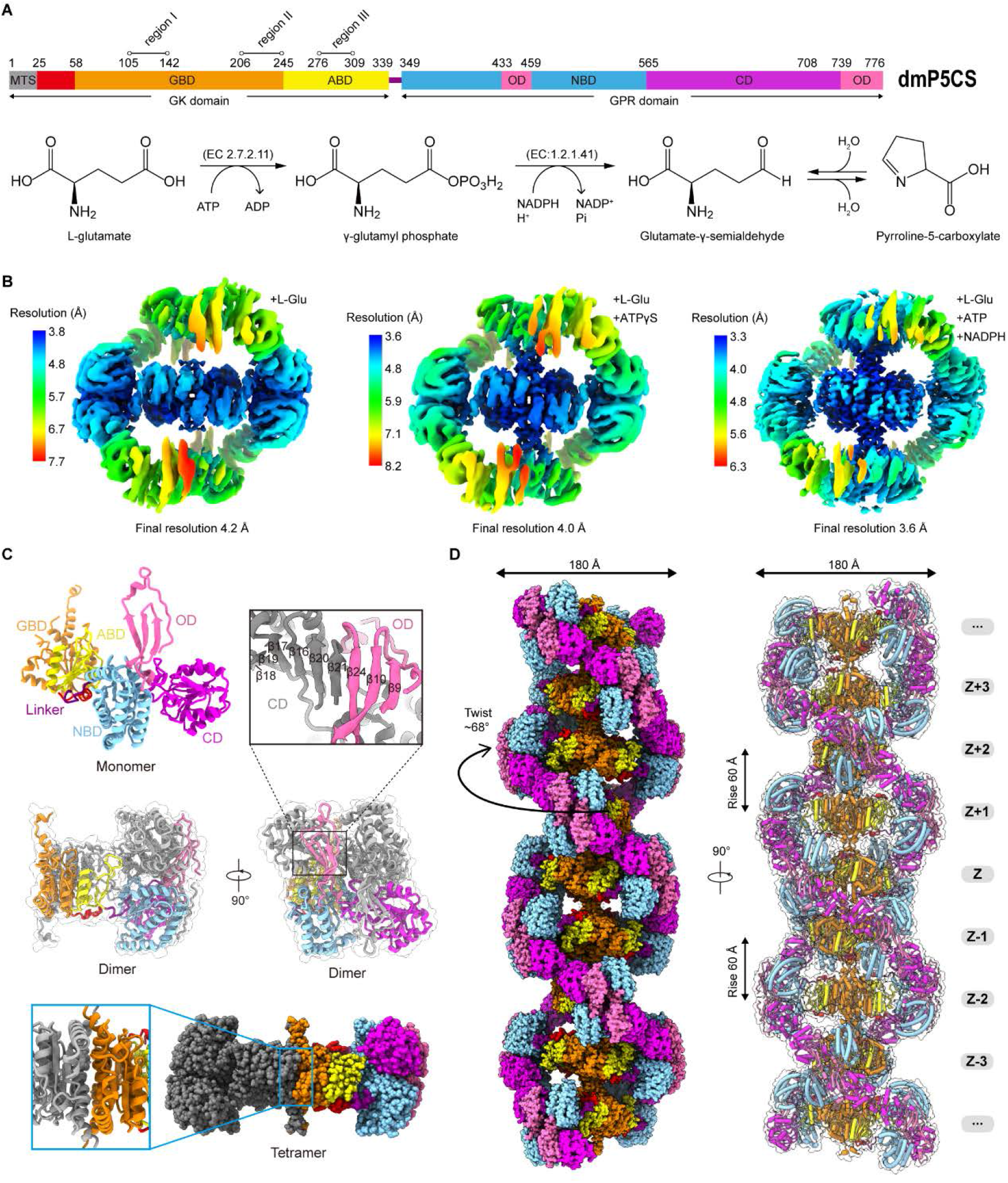
Bifunctional enzyme properties and cryo-EM analysis of P5CS filament. (**A**) Domain organization of *Drosophila melanogaster* P5CS, that consists of two domains, N-terminal glutamate kinase (GK) domain and C-terminal glutamyl phosphate reductase (GPR) domain. Putative mitochondrial targeting sequence (MTS) labelled in grey, the glutamate binding domain (GBD) and the ATP binding domain (ABD) of GK domain are respectively shown in orange and chartreuse, the NADPH binding domain (NBD), the catalytic domain (CD) and the oligomerization domain (OD) of GPR domain are shown in cyan, purple, and pink, respectively. Bifunctional P5CS enzyme catalytic reaction and residue numbers for domain boundaries are shown. (**B**-**D**) Single particle analysis for 3D reconstruction of P5CS filament, three cryo-EM maps of P5CS^Glu^ filament, P5CS^Glu/ATPγS^ filament and P5CS^Mix^ filament, which formed in different substrate environment incubation, colored by local-resolution estimations. (**E**) The structures of P5CS monomer, color codes for P5CS models are indicated. (**F**) The P5CS dimer. Two monomers (grey or color coded by domain) interact via GPR domain hairpins contact. (**G**) The P5CS tetramer (sphere representation) is formed via GK domain interaction (cartoon representation) between two P5CS dimer (grey or color coded by domain). (**H**) The sphere and cartoon representation of P5CS filament. P5CS filament are modelled by the cryo-EM map, rotated view is shown in the right panel, rise, twist, and width are indicated.

Subsequently, P5CS protein was incubated with different combinations of substrates for the preparation of samples for cryo-electron microscopy (Figure S1D-G). Long and flexible filaments of P5CS were observed under all conditions (Movie S1 and S2). Filaments in three conditions with glutamate (P5CS^Glu^), glutamate and ATPγS (P5CS^Glu/ATPγS^), and glutamate, ATP and NADPH (P5CS^Mix^), were imaged in cryo-EM for single particle analysis (SPA). After 3D classification and 3D reconstruction, the electron density maps of the P5CS^Glu^, P5CS^Glu/ATPγS^ and the P5CS^Mix^ filaments achieved at a resolution of 4.0 Å, 4.2 Å and 3.6 Å, respectively (Figure 1B-D, S2 and S3). Using separate focused refinement strategy, we obtained multiple different conformational states of GK domain tetramer structures (3.1 to 3.5 Å) and GPR domain dimer structures (3.6 to 4.2 Å). The cryo-EM data and model refinement statistics are provided in Table 1. The N-terminus (Residues 1–44) is invisible in our maps, and three disordered fragments in region I, II and III in GK domain are depicted by dashed lines in our models.

One P5CS monomer can be roughly divided into five distinct subdomains: the glutamate binding domain (GBD) and the ATP binding domain (ABD) at the GK domain; the NADPH binding domain (NBD), the catalytic domain (CD) and the oligomerization domain (OD) at the GPR domain (Figure 1A, E). In the model, two P5CS monomers dimerizes through the interaction between their GPR domains, where the β21 at the CD interacts with the β24 at the OD of another monomer (Figure 1F). This interaction connects two groups of hairpins and maintains the homodimer structure by a hydrogen bond network. Two P5CS dimers further assemble a compact tetramer through the interaction at the GK domains, becoming the building blocks for the filament (Figure 1G).

We examined the P5CS filament structures in these states shows that the characteristics of each helical P5CS filament is highly similar with the binding of different ligands. We chose the P5CS^Mix^ filament to display structural details (Figure 1H). In the spiral P5CS filament structure, GK domains tetramers serve as the core of the filament, and the GPR domains dimer forms a left-handed double helix structure around the central axis. The overall diameter of P5CS filaments in all three states is 180 Å, the helical twist is 68°, and the helical rise is 60 Å (Figure 1H).

### Structural comparison with ligand-bound GK domain

The structures of *Drosophila* P5CS GK domain and *E. coli* GK are highly conserved. The sequence and structural alignments suggest that their secondary structures are similar as both of them exhibit a sandwich-like α3β8α4 topological folding and two disordered loops at region II and region III. These disordered loops are located near the binding-pocket of the GK domain. An opening loop in region II was observed in P5CS^Glu/ATPγS^ and P5CS^mix^ structures, while closure was found in the P5CS^Glu^ structure (Figure 2A, B). We speculated that this fragment may participate in the regulation of the binding of ligands.

**Figure 2.**
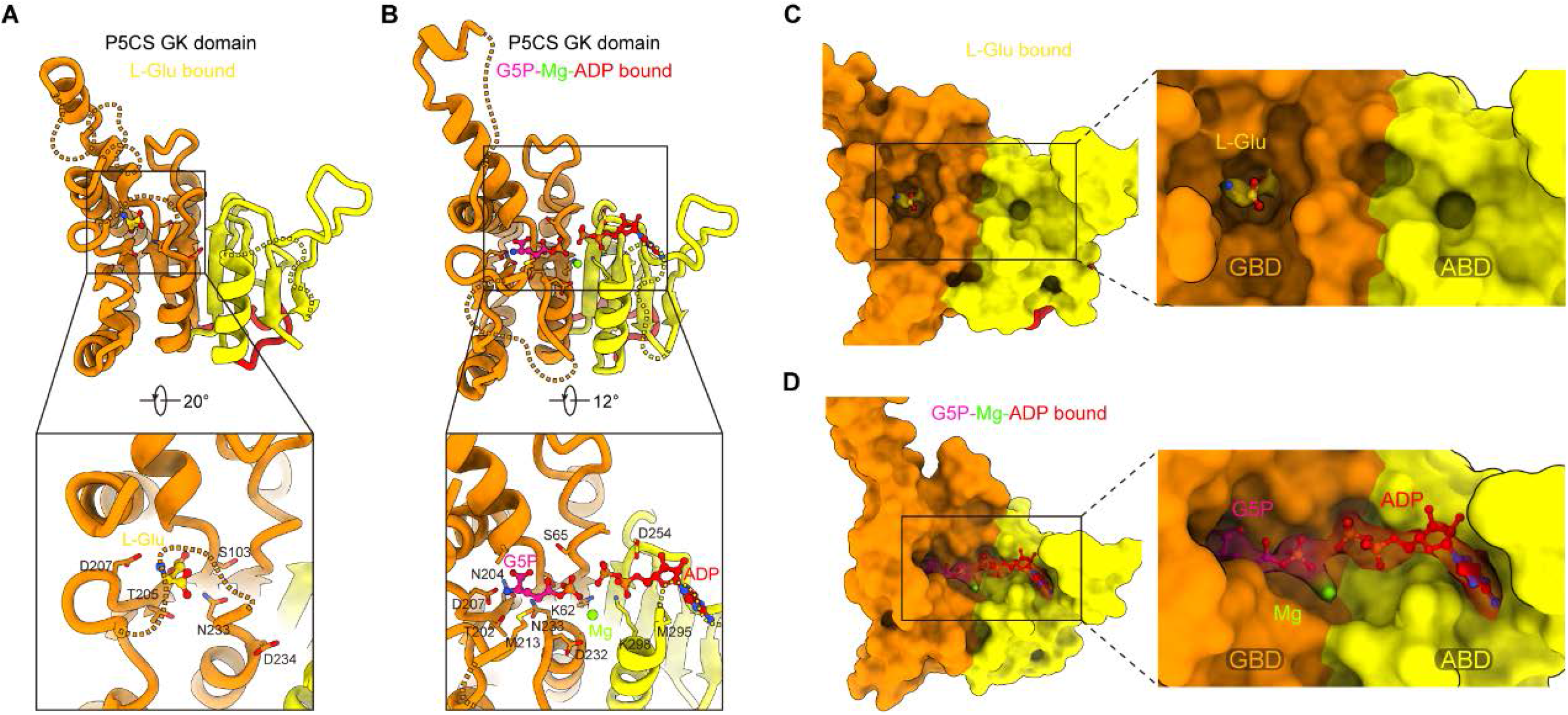
Conformational changes in GK domain binding pocket. (**A**) GK domain of the P5CS^Glu^ filament, with L-glutamate shown as sticks with yellow carbons. (**B**) GK domain of the P5CS^Mix^ filament, with G5P, Mg^+^ and ADP shown as sticks with pink, green and red carbons, respectively. (**C**-**D**) GK domain model surface representation showing the conformation of binding pocket in P5CS^Glu^ filament or in P5CS^Mix^ filament. The cryo-EM density of a bound glutamate molecule in (**C**), a bound complex of G5P, Mg^+^ and ADP in (**D**).

Three conformations of the GK domain with the binding of different ligands were revealed clearly in our models. In the GK domain, a valley-like pocket is located between the GBD and the ABD, providing the binding sites for L-glutamate, ATP or their derivatives (Figure 2C, D). In the P5CS^Glu^ filament, L-glutamate is bound in a vertical way, the closure loop in region II make glutamate relatively enclosed in the GBD (Figure 2A, C). In contrast, in the P5CS^Mix^ filament, the glutamate at the binding site is converted into the intermediate G5P, and the phosphate donor ATP becomes an ADP at the ABD with the association with a Mg^2+^ (Figure 2B, D). Meanwhile, the loop shifts away from the top of the binding pocket (Figure 3A), and residue M213 interacts with G5P in this conformation.

**Figure 3.**
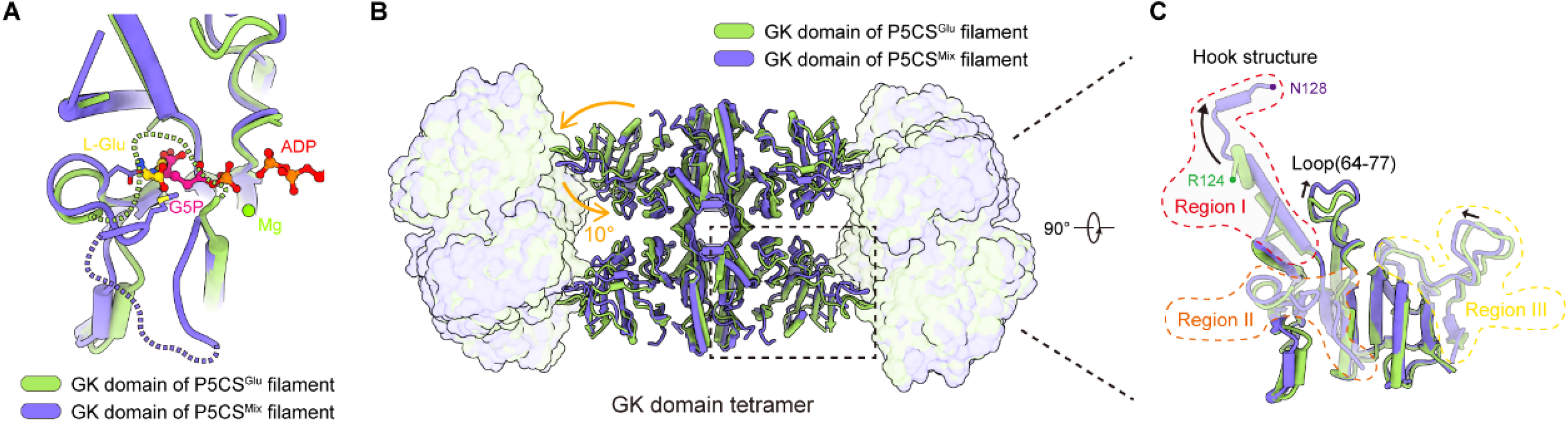
Structural comparison of two types of GK domain. (**A**) Comparison of the GBD of GK domain in the P5CS^Glu^ filament (green) with the P5CS^Mix^ filament (purple), corresponding to the loop in region II at closed conformation in P5CS^Glu^ filament and open conformation in P5CS^Mix^ filament. (**B**) Superimposition of GK domain tetramer in the P5CS^glu^ filament (green) with the P5CS^Mix^ filament (purple). Transitions from glutamate bound conformation to G5P-Mg-ADP bound conformation are shown as curved arrows, indicating GK domain conformational changes in P5CS filament. (**C**) Comparison of one protomer of GK domain tetramer in P5CS^Glu^ filament (green) and P5CS^Mix^ filament (purple), the disorder fragments are hided in our model.

We compared structures of the GK domain in the P5CS^Glu^ and P5CS^Mix^ filament to investigate conformational changes involved in the catalytic reaction. (Figure 3B). First, the GK domain of each protomer rotates approximately 10° around its central axis (Figure 3B), causing the horizontal compression of the GK domain dimer. Second, the helix-helix structure (Residues 105-113, 115-124) at the region I of P5CS^Glu^ transforms into a helix-loop-helix structure (Residues 105-119, 120-122, 123-128) in the P5CS^mix^ (Figure 3C). This helix-loop-helix structure is referred to as the “hook” structure. The transformation of the hook structure results in new contact sites between neighbor tetramers in the vertical direction, which can be evidenced by a rigid density in our map (Figure 1B-D). On the other hand, we also noticed a loop (Residues 64-77) shifts greatly by approximately 3 Å (Figure 3C). However, the function of this conformational change is not clear. The structure of GK domain in the P5CS^Glu/ATPγS^ filament largely overlaps with the P5CS^Mix^, suggesting given conformational changes are mainly regulated by the binding of ATP/ADP.

Together, by comparing the structures of GK domain with different ligands, we reveal conformational changes during the phosphorylation of glutamate. The binding pocket transforms into a relatively loose conformation upon the binding of ATP/ADP. In this conformation, the glutamate and ATP are extended towards each other, creating an active state for the reaction.

### Open and closed conformation of GPR domain

On the basis of P5CS structures, we display four different binding modes (Figure 4A-D) of GPR domain. GPR domain is in an unliganded state in the P5CS^Glu^ filament (Figure 4A). In the P5CS^Glu/ATPγS^, however, we observed the density of a G5P at the CD active site (Figure 4B). The unexpected presence of this G5P could be due to the contamination of ATP in the commercial ATPγS, of which the purity is >80%. With this model, the binding mode of this G5P is clearly resolved. This state is referred to as the G5P-binging state. In focus refinement of GPR dimmer structures of P5CS^Mix^ filament, we determined two other states of the GPR domain (Figure 4C, D). One is NADP(H)-binding state, which NADP(H) is present at the NBD. Another one is NADP(H)-released state, and cofactor binding site is empty.

**Figure 4.**
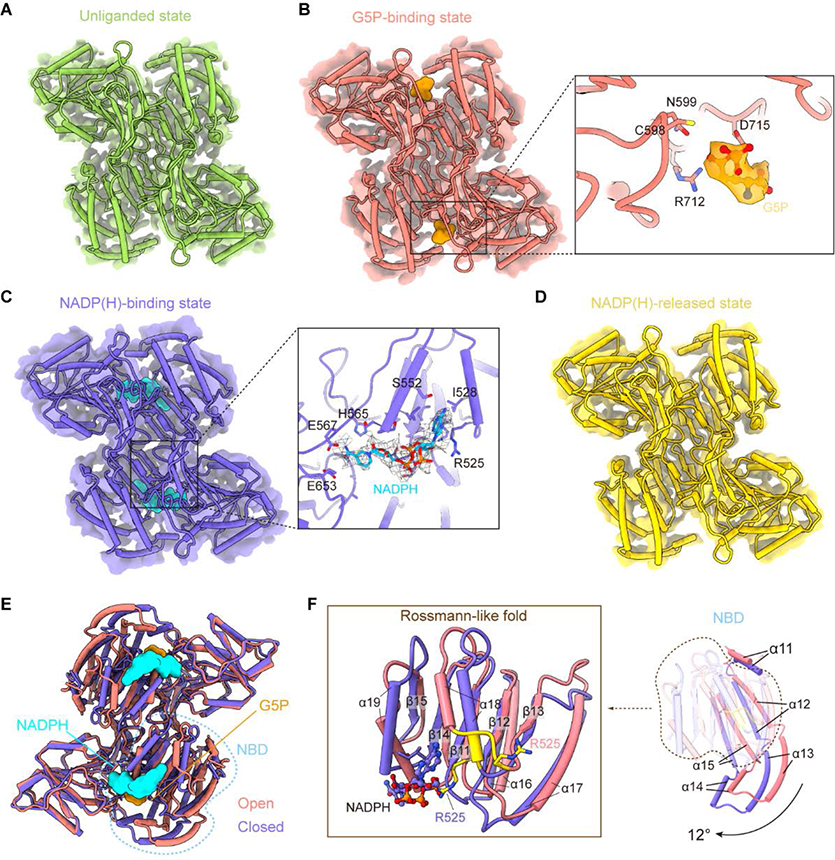
GPR domain ligand-bound mode and its conformation. (**A**) The cryo-EM density of GPR dimer structure and cartoon model are representative as unliganded state in P5CS^Glu^ filament (green). (**B**) GPR dimer structure of G5P-binding state in P5CS^Glu/ATPγS^ filament (coral). The conformation of G5P-binding pocket, with G5P (orange) is shown as sticks carbons. (**C**) GPR dimer structure of NADP(H)-binding state in P5CS^Mix^ filament (purple). The conformation of NADP(H)-binding pocket, with NADPH (cyan) is shown as sticks carbons. (**D**) GPR dimer structure of NADP(H)-released state in P5CS^Mix^ filament (yellow). (**E**) Structural differences in the G5P-binding state (coral) and NADP(H)-binding state (purple) of GPR domain. Ligands are colored as in (**B** and **C**). (**F**) Superimposition of the either NBD or the Rossmann-fold of GPR domain at G5P-binding state and NADP(H)-binding state using single protomer.

Comparison of the conformation of four states of described above shows that the overall structures of GPR domain are similar except the NADP(H)-binding state. Interestingly, while the structures of CD and the OD are generally consistent in all four states, the NBD of the GPR domain in the NADP(H)-binding state greatly differs from others (Figure 4E). The NBD contains consecutive alternating α-helices and β-strands (α2-β5-α2) architecture, which is known as the Rossmann-like fold for dinucleotide-binding(*18, 19*). By superimposing this Rossmann-like fold and the entire NBD, we determined conformational changes between single GPR domain at the G5P-binding and NADP(H)-binding state (Figure 4F). Upon NADP(H) binding, the residue R525 interacts with its adenine moiety. This interaction transforms the ^525^REE^527^ loop into an ordered structure that extends the α18-helix (Figure 4F) in Rossmann-like fold. Meanwhile, the entire NBD rotates approximately 12° about the β9-β10 motif of the OD and slides towards the CD (Figure 4F; Movie S3). This transformation contributes to bring the nicotinamide ring of the NADP(H) approaching to C598 of the CD. The C598 of CD is considered as a crucial amino acid for catalysis in ALDH family(*20*). Accordingly, these results indicate that such conformational changes are triggered by the binding of NADPH to initiate the transfer of the hydride ion from NADPH to the intermediate G5P.

We suggested that unliganded, G5P-binding and NADP(H)-released states represent as open conformation, and the NADP(H)-binding state represents as closed conformation. We propose that the P5CS filament may accommodate the GPR domain in both conformations, and the repetitive transformation between these two conformations is essential for the catalytic reaction.

### Filamentation regulates the enzyme activity

In P5CS^Glu^, P5CS^Glu/ATPγS^ and P5CS^Mix^ filaments, the neighboring GPR domain dimers interact with each other to form a spiral structure. The CH/Pi interaction(*21*) formed between F642-P644 of the contact loops in adjacent GPR domains is proposed to be responsible for the filamentation (Figure 5A, B). In P5CS^Glu/ATPγS^ and P5CS^Mix^ filaments, additional interface locks adjacent GK domain tetramers within filament by hook structures pairs. The hook structure extrudes from GK domains and towards their counter parts in the adjacent GK domain tetramers (Figure 5A, C), forming strongly contacts via hydrogen bonds (M119-R124, L121-M123) and salt bridges (E116-R124). These interactions at different contact sites in the P5CS filament stabilize the double helix architecture and its inner core, respectively.

**Figure 5.**
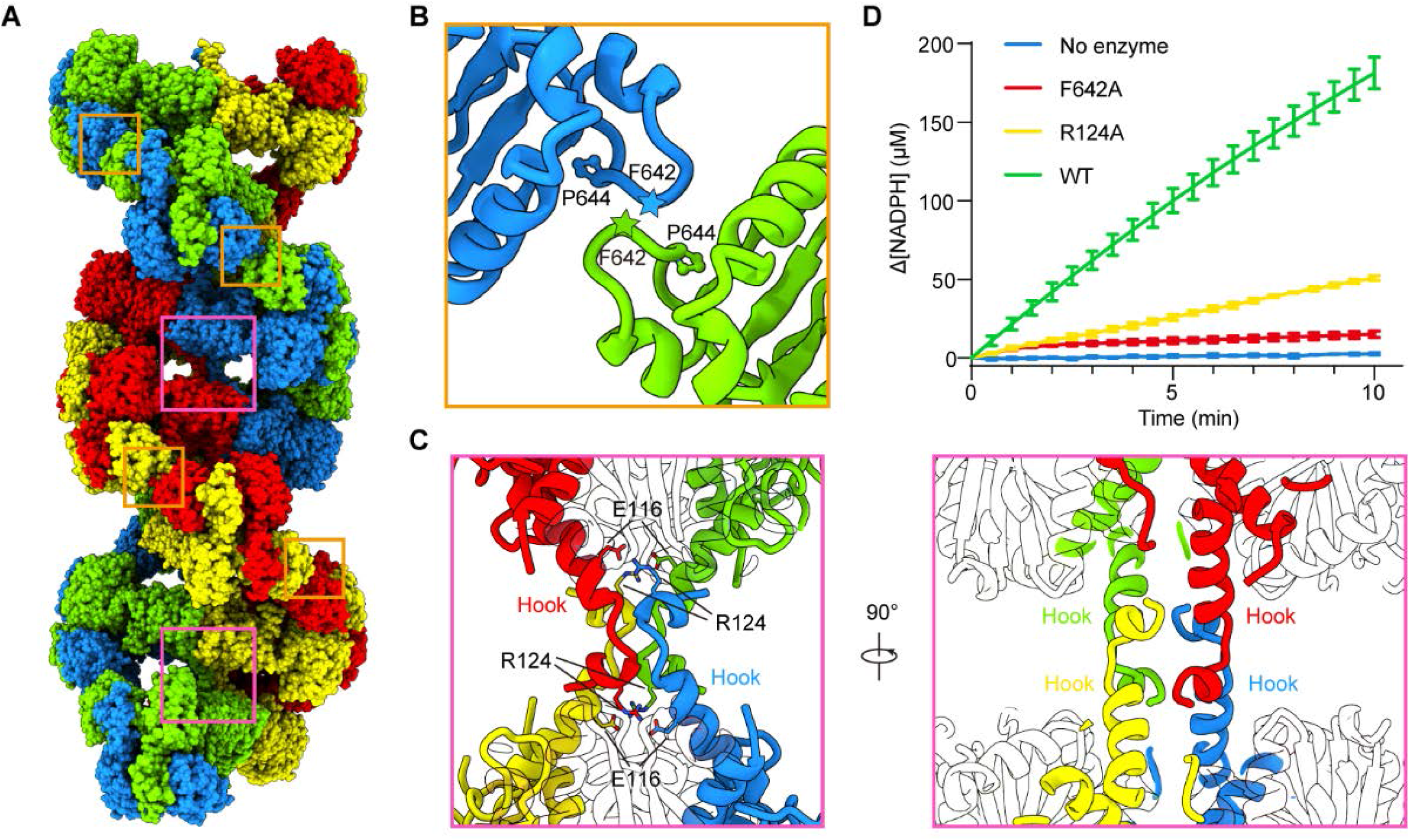
Assembly and interaction surfaces of the P5CS filament. (**A**) P5CS filament assembly interface, the four P5CS monomers in one layer are colored in red, yellow, blue and green. **(B**) Interaction between two adjacent GPR domain dimers, residues F642 located at loop which interacts with P644 from another neighboring GPR domain dimer. (**C**) Model for hook structure interaction. (**D**) Enzyme activity analysis to examine P5CS wild-type or mutant proteins. All of the experiments were biologically replicated three times (n = 3, mean ± s.d.).

To understand the functions of P5CS filaments, we generated two mutants, R124A and F642A, which are predicted to abrogate the tetramer-tetramer contact sites of GK domains and GPR domains, respectively. Negative stain of mutant P5CS shows that the P5CS^F642A^ mutant proteins do not assemble into filament in any condition (Figure S4A). These results indicate that the interaction at the GPR domain interface is crucial for P5CS filamentation. In contrast, the P5CS^R124A^ mutant proteins formed long filaments in the APO state as well as in the condition with L-glutamate (Figure S4B). However, when ATP was supplemented to the glutamate-bound P5CS^R124A^ filaments, the depolymerization of those filaments were observed. When being incubated with all substrates, the P5CS^R124A^ mutant formed shorter filaments in comparison with P5CS^WT^ (Figure S4B). Based on our models, the binding of ATP is predicted to initiate the reaction at the GK domain, and also to trigger the creation of hook structures (Figure 5C). We propose that the interaction of hook structure pairs is necessary for stabilizing the filament during the transformation from the P5CS^Glu^ filament to the P5CS^Glu/ ATPγS^ filament or P5CS^Mix^ filament.

We subsequently analyse the activity of wildtype P5CS and two mutants. Comparing with wildtype P5CS, R124A and F642A mutants exhibited a dramatically compromised activity (Figure 5D), indicating that the integrity of filamentation is essential for the catalytic reactions. Since the mutated residues are not supposed to directly interfere the active sites, the function of P5CS filament is suggested as to couple the reactions catalyzed at two domains. First, the intermediate G5P is instability and the distance between two reaction centers is about 60 Å (Figure S5), so the filament of P5CS may exhibit a scaffold architecture that stabilize the relative position of GK and GPR domain or electrostatic substrate channeling between two domains. Second, P5CS filamentation may create a half-opened chamber, as the active sites of P5CS are located at the inner part of the filament, and GK domain is catalytically fast compared to GPR domain. Therefore, the unstable intermediates G5P are abundant within the filament, this microenvironment will reduce the release of G5P into the solvent, thereby facilitating the rate-limiting reaction at the GPR domain. Thus, the disruption of P5CS filament may result in uncoupled catalytic reactions of bifunctional P5CS and a reduced activity.

### Mechanisms of P5CS filament catalysis

In the proposed model, spontaneous filamentation and elongation of P5CS is importantly associated with the binding of glutamate. The loose pocket at the GK domain is subsequently bound by ATP and extrudes the hook structure along with the phosphorylation of the glutamate. When the products of the GK domain dissociate from the pocket, the G5P is trapped within the filament and further captured by the GPR domain. Next, NADPH binds to the GPR domain, the conformational change brings the NADPH towards the catalytic residue C598, becoming the close conformation and facilitating the reaction. After the reaction is accomplished, NADP^+^ and GSA will be released and the GPR domain returns to its open conformation (Figure 6). This working mechanism suggests that GK and GPR domain undergo continuous conformational transitions during catalysis, making it a dynamic filament.

**Figure 6.**
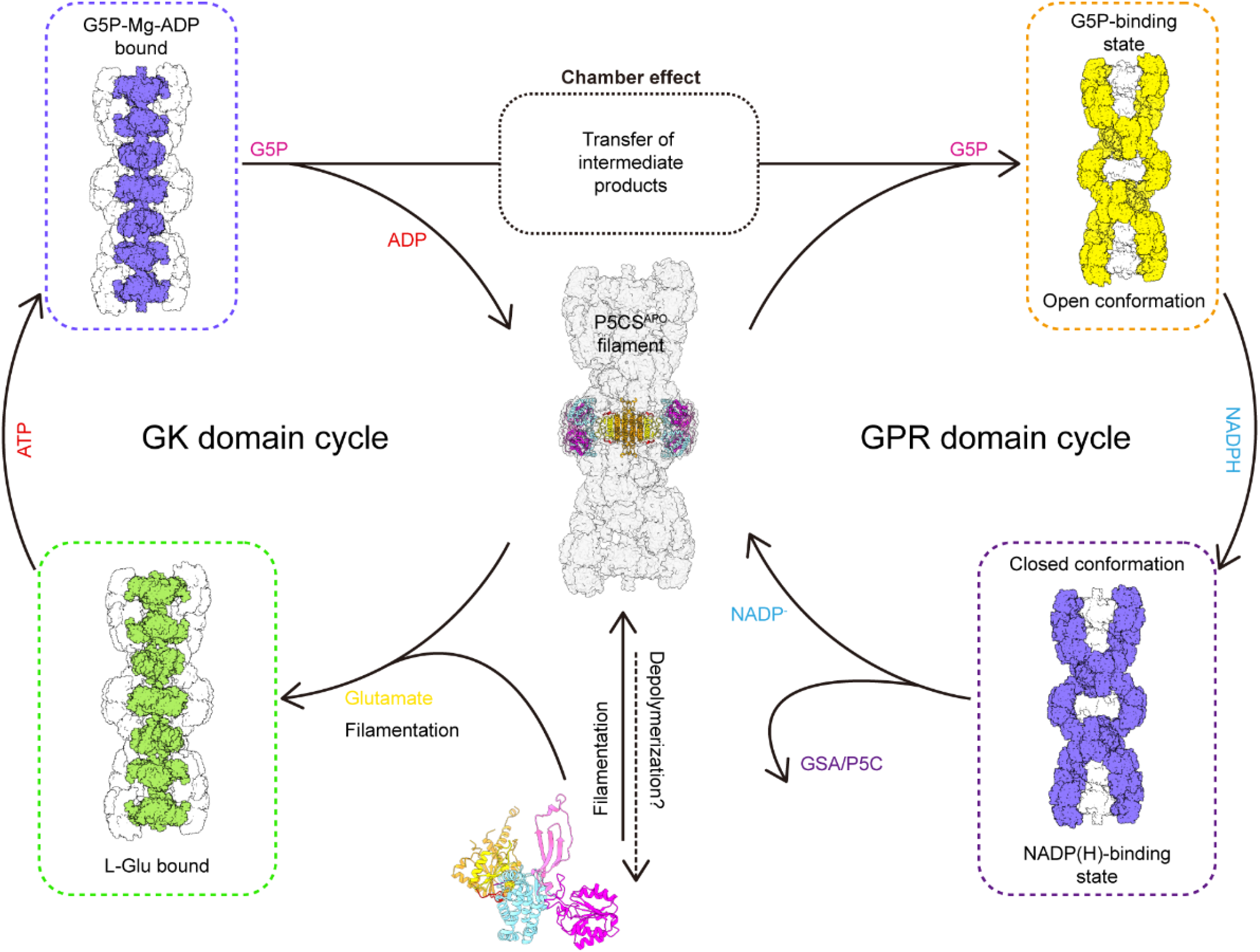
Model of P5CS filament structural transitions during GSA/P5C synthesis. The P5CS molecule polymerizes into filaments at apo state or after binding with the glutamate. Upon ATP binding, GK domain initiates glutamate phosphorylation. The product leaves the pocket and GK domain subsequently repeat reaction cycle (left). Unstable G5P will be transported through the half-open chamber inside the filament, and captured by GPR domain. NADPH binding to GPR domain that transform to open conformation, which enables NADPH approach to catalytic site and completes reductive dephosphorylation of G5P. The GSA/P5C will be released and the GPR domain returns to unliganded state with open conformation, GPR domain begins the next cycle.

Recently, increasing reports about hyperammonaemia, neurocutaneous syndromes and motor neuron syndrome caused by mutations on human P5CS gene (*ALDH18A1*) have been published. Such mutations may result in the loss of P5CS function in various degrees. *Drosophila* P5CS residue R124 in the region I of the GK domain, which correspond to R138 of human P5CS, is highly conserved among different eukaryotes (Figure S6). However, 17 residues in the region I, including the hook structure, is absent in *E. coli* GK. Pathogenic mutations of R138 in human P5CS, which is proven to be the cause of autosomal-dominant cutis laxa, have been demonstrated with a decreased activity and a dispersed distribution in mitochondria(*22, 23*). Herein, the structural basis reveals that R138 mutation on human P5CS could abrogate the interaction between hook structures of GK domains, destabilizing the filament and the coupling of reactions.

In summary, our cryo-EM structures of *Drosophila* P5CS filament present the assembly mode of P5CS protein, and provide insights into the molecule basis of the coupling of GK and GPR domains in the filamentous structure.

## Methods

### P5CS Protein purification

The full-length *Drosophila melanogaster* P5CS gene were cloned into a modified pET28a vector with a 6×His–SUMO tag fused at the N terminus, the fusion proteins were expressed in *E. coli* Transetta (DE3) cells overnight at 16°C after induction with 0.1mM IPTG at OD600 range of 0.6∼0.8. The remainder of purification was performed at 4 °C. The harvested cells were sonicated under ice and purified by Ni-NTA agarose beads (Qiagen) in lysis buffer (50 mM Tris-HCl pH 8.0, 500 mM NaCl, 10% glycerol, 20 mM imidazole, 1 mM PMSF, 5 mM β-mercaptoethanol, 5 mM benzamidine, 2 μg/ml leupeptin and 2 μg/ml pepstatin). After in-column washing with lysis buffer, the proteins were eluted with elution buffer (50 mM Tris-HCl pH 8.0, 500 mM NaCl, 250 mM imidazole, 5 mM β-mercaptoethanol), peak fractions were treated with SUMO protease for 1 h at 8 °C. The P5CS proteins were further purified through HiLoad 16/600 Superdex 200 pg gel-filtration chromatography (GE Healthcare) in column buffer (25 mM HEPES pH 7.5 and 100 mM KCl), peak fractions were collected, concentrated and stored at −80°C before to use.

### Enzyme activity assays

The full-length wild type or mutant P5CS activity was determined in the reaction buffer containing 25 mM HEPES pH 7.5, and 10 mM L-glutamate, with added 20 mM MgCl2, 10 mM ATP, and 0.5 mM NADPH used to initiate the reaction, then the reaction was monitored at 37 °C in a MD-SpectraMax i3 plate reader and absorbance at 340 nm was measured every 20 s for 10 min (one experiment, n = 3 each). The NADPH concentration was converted from A340 with the standard curve determined at the same experiment.

### Negative staining

Wild-type or mutation P5CS proteins were mixed with different substrate conditions. In brief, the final concentration was as follows: 25 mM HEPES pH 7.5, 100 mM KCl, 10mM MgSO4, 100 mM L-glutamate, 10 mM ATP and 0.5 mM NADPH. The prepared protein samples were applied to glow-discharged carbon-coated EM grids (400 mech, zjky), and stained with 1% uranyl acetate. Negative-stain EM grids were photographed on a Tecnai Spirit G21 microscope (FEI).

### Cryo-EM grid preparation and data collection

For cryo-EM, purified full-length P5CS was diluted to approximately 2 μM and dissolved in buffer contains 25 mM HEPES pH 7.5, 100 mM KCl, 10mM MgSO4, and incubated with 20mM L-glutamate for the P5CS^Glu^ filament preparation. The P5CS^Glu/ATPγS^ filament additionally added with 0.5mM ATPγS compare to the P5CS^Glu^ filament. For the P5CS^Mix^ filament, P5CS proteins (2 μM) incubated with 100 mM KCl, 10mM MgSO4, 20mM L-glutamate, 2 mM ATP and 0.5 mM NADPH. All the samples incubated for 1h on ice before vitrification. The P5CS filament samples were placed on H2/O2 glow-discharged holey carbon grids (Quantifoil Cu 300 mesh, R1.2/1.3) or amorphous alloy film (No. M024-Au300-R12/13). Then Grids were immediately blotted for 3.0 s and plunge-frozen in liquid ethane cooled by liquid nitrogen using Vitrobot (Thermo Fisher) at 4 °C and with 100% humidity. Image were collected on Titan Krios G3 (FEI) equipped with a K3 Summit direct electron detector (Gatan) in direct in counting super-resolution mode at 300 kV with a total dose of 72 e^−^/Å^2^, subdivided into 50 frames in 4 s exposure using SerialEM. The images were recorded at a nominal magnification of 22,500× and a calibrated pixel size of 1.06 Å, with a defocus ranging from 0.8 to 2.5 μm.

### Image processing and 3D reconstruction

The whole image analysis was performed with RELION. We use MotionCor2 and CTFFIND4 via RELION GUI to pre-process the image, movie frames were aligned and the contrast transfer function (CTF) parameters were estimated in this progress. After manual selection, there are 4933 images for P5CS^Glu^ dataset, 6408 images for P5CS^Glu/ATPγS^ dataset, 10566 images for P5CS^Mix^ dataset left for the further processing. For the flexibility of P5CS filaments, single particle analysis was carried out in our reconstructions and no helical symmetry was implied in the whole process. Reference free particle picking built in RELION3 was performed. This process provides 1994786 particles for P5CS^Glu^, 2024372 particles for P5CS^Glu/ATPγS^ and 8027582 particles for P5CS^Mix^. At first, particles were extracted binning 2 or 3 times for the fast 2D classifications. Datasets were cleaned with several rounds of 2D classification and the bin factors were gradually reduced to 1 at the same time. After 3D classifications with C1 symmetry applied, several classes were selected to do finer 3D classifications with D2 symmetry. Classes with intact structure were retained for 3D refinement with D2 symmetry. For the 3D refinement, 432746, 327841, and 1412498 particles were used for each dataset. And map include three P5CS tetramer layers were obtained. The relative motion between GK and GPR limited the refinement at high resolution, so we use partition reconstruction strategy to improvement the resolution for both GK and GPR domain. For GK domain, we use continued local refinement to improve the resolution with a mask focus the middle layer GK. Then the Ctf-refinement and Bayesian polishing were performed for the remained particles and improve the resolution to 4.2 Å, 4.1 Å, and 3.6 Å for 3-layer P5CS maps and 3.5 Å, 3.4 Å, and 3.1 Å for GK maps. For GPR domain, particles were expended symmetry for the 3D classification without alignment. Several classes with intact structure were selected and oriented, symmetry collapse was done at the same time. Then 3D classifications and refinements with C2 symmetry were performed. For the P5CS^Mix^, two different states of GPR were captured. Finally, we got 286291. 348804, 193482, and 233624 particles to construct maps for GPR domain with 3.6 Å, 4.2 Å, 4.3 Å and 4.2 Å resolution. LocalRes was used to estimate the local resolution of our map.

### Model building refinement and validation

Based on our map with near atomics resolution, the model of GK and GPR were generated with focused refinement map for different state. The initial model of GK domain and GPR domain were generated via swiss-model regarding 4Q1T and 2h5g as reference separately. Manual adjustment and building the missing regions were done in Coot. Real space refinements were performed by Phenix. The full-length P5CS model are linked using corresponding GK and GPR structures, the linker was generated in the Coot and refined via Phenix. Figures and movies were generated by UCSF Chimera and ChimeraX.

## Acknowledgements

We thank Zhi-Jie Liu and Suwen Zhao for helpful discussions. The EM data were collected at the ShanghaiTech Cryo-EM Imaging Facility. We thank the Molecular and Cell Biology Core Facility (MCBCF) at the School of Life Science and Technology, ShanghaiTech University for providing technical support.

## Supplementary information

**Figure S1.**
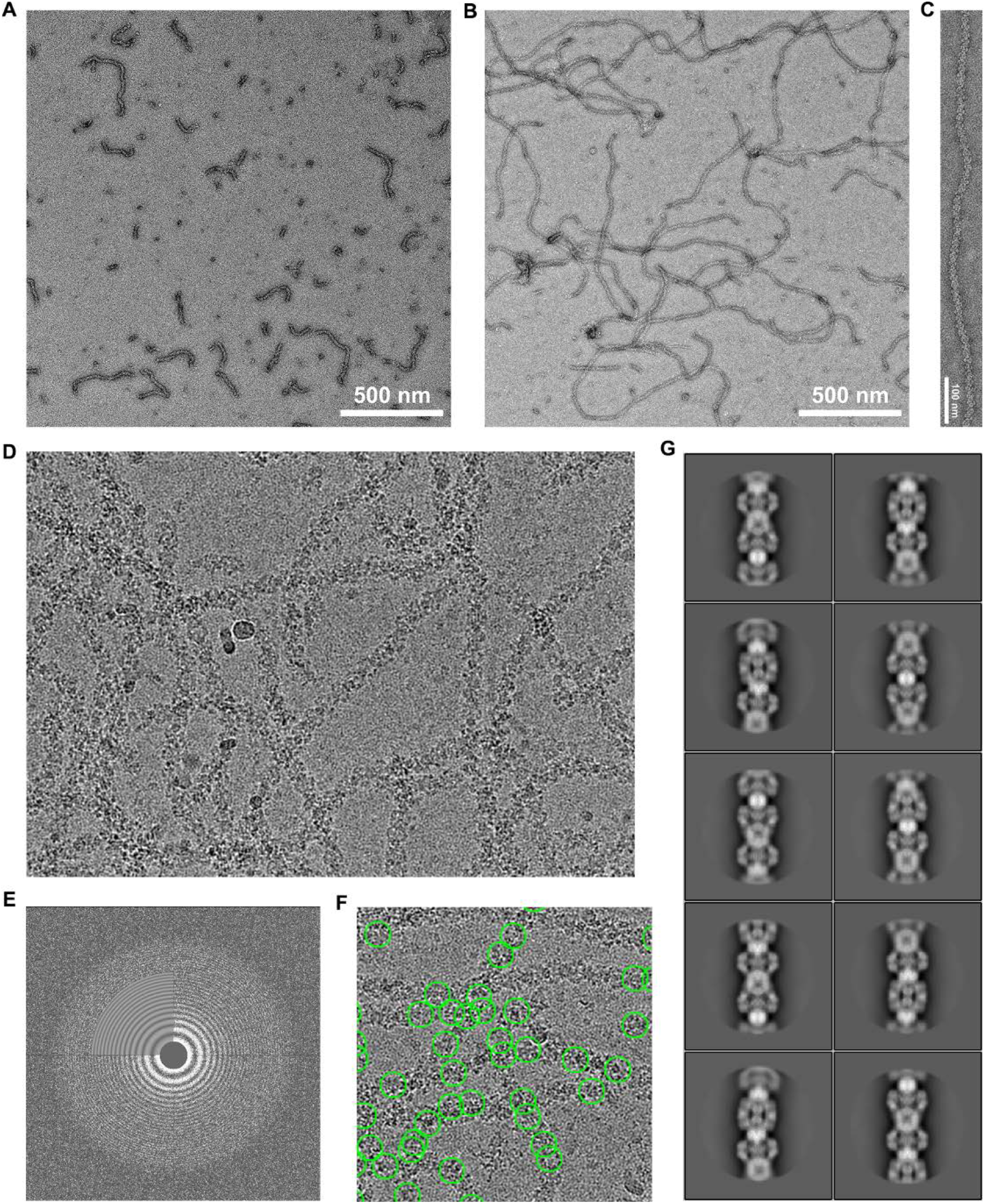
Substrates can significantly extend the P5CS filament. (**A**) Negative stain electron microscopy micrograph of P5CS protein in APO state, we find that the P5CS protein in the apo state can self-assemble into filaments of various lengths. (**B**) When substrate (KCl, MgSO4, ATP, NADPH and glutamate) were added to the reaction which can promote the extension of P5CS protein filament. (**C**) Representative negative stain electron microscopy micrographs of a single P5CS filament in substrate state. (**D**) A representative cryo-EM micrograph of P5CS filament. (**E**) The power spectrum of the micrograph. (**F**) The green circles representing single particles of the picked P5CS filament. (**G**) Representative 2D class averages of P5CS filament, several classes of P5CS filament particles with less curvature were selected.

**Figure S2.**
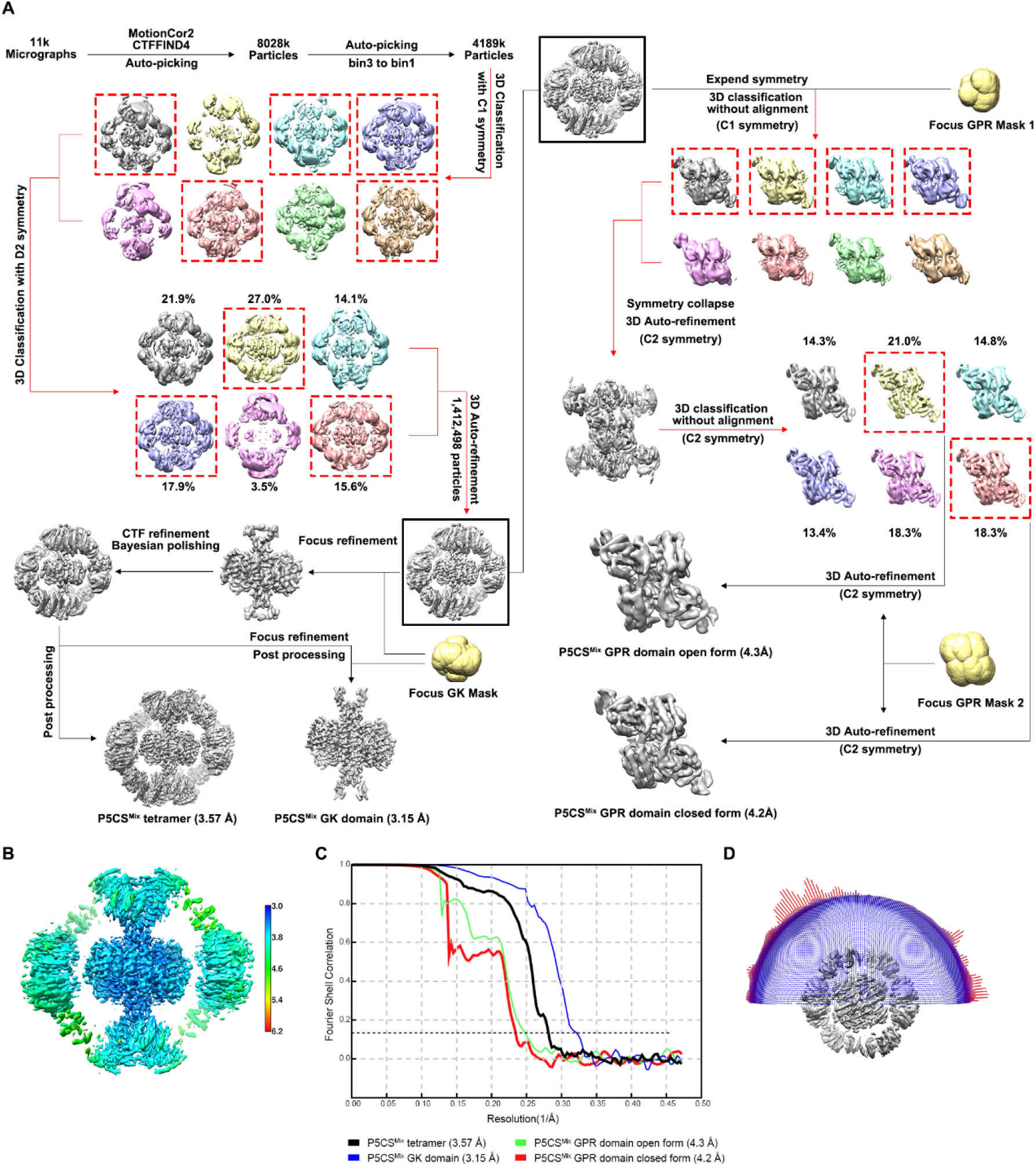
cryo-EM analysis of P5CS^Mix^ filament. (**A**) Flow chart for the cryo-EM reconstruction of P5CS^Mix^ filament, additional detailed procedures were described in Methods. (**B**) Local-resolution map of the P5CS^Mix^ tetramer. (**C**) The gold-standard Fourier shell correlation curves for four refined maps. (**D**) Angular distribution of the particles used for the final reconstruction of P5CS^Mix^ filament.

**Figure S3.**
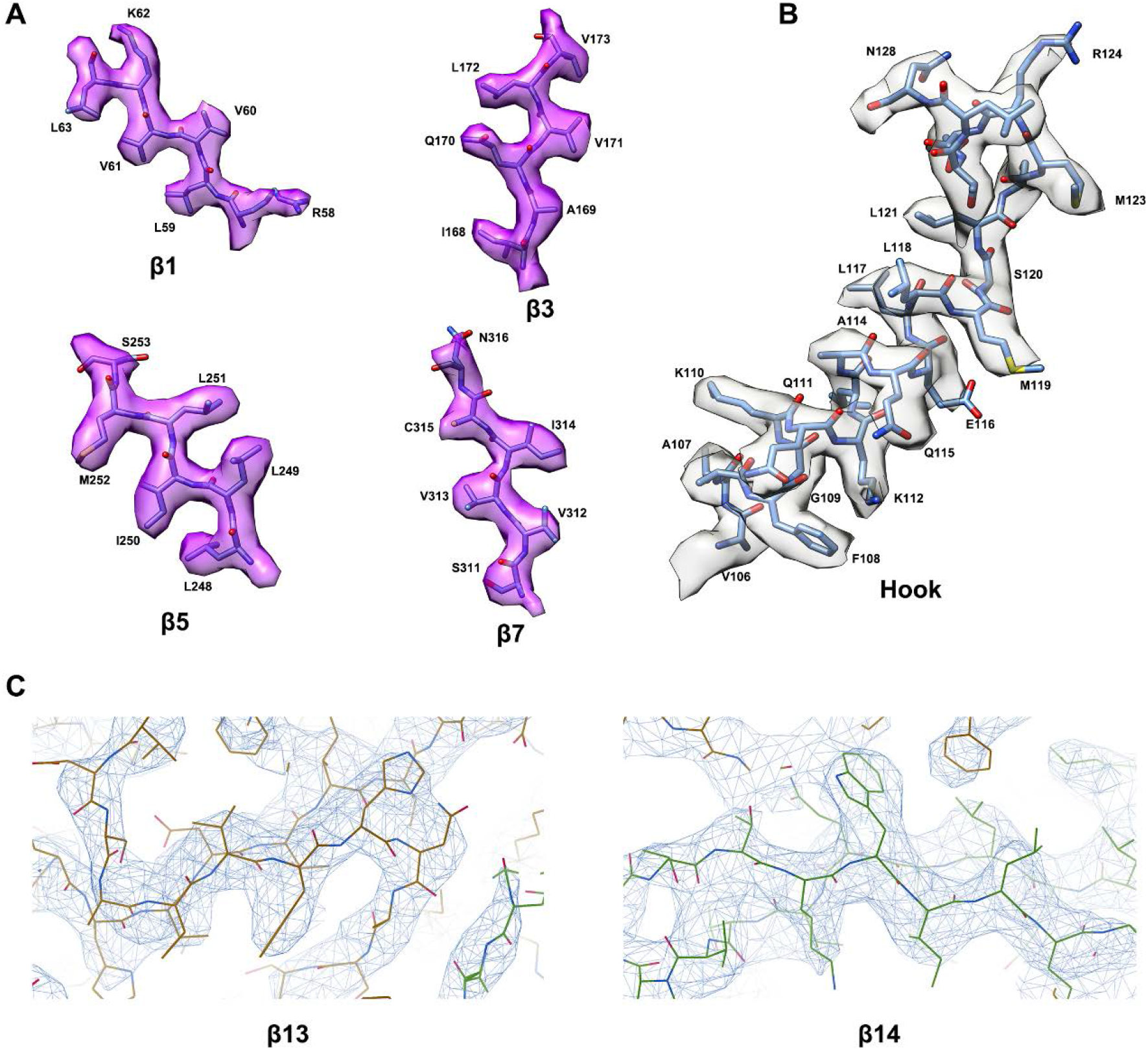
Representative cryo-EM map. Representative regions of the P5CS protein model, superimposed cryo-EM map. (**A**) Atomic model of P5CS β-sheets. (**B**) Atomic model of the hook structure α-helices. (**C**) Representative cryo-EM density of β13 and β14 is displayed at 4.5 σ contour level, panel C is generated by COOT.

**Figure S4.**
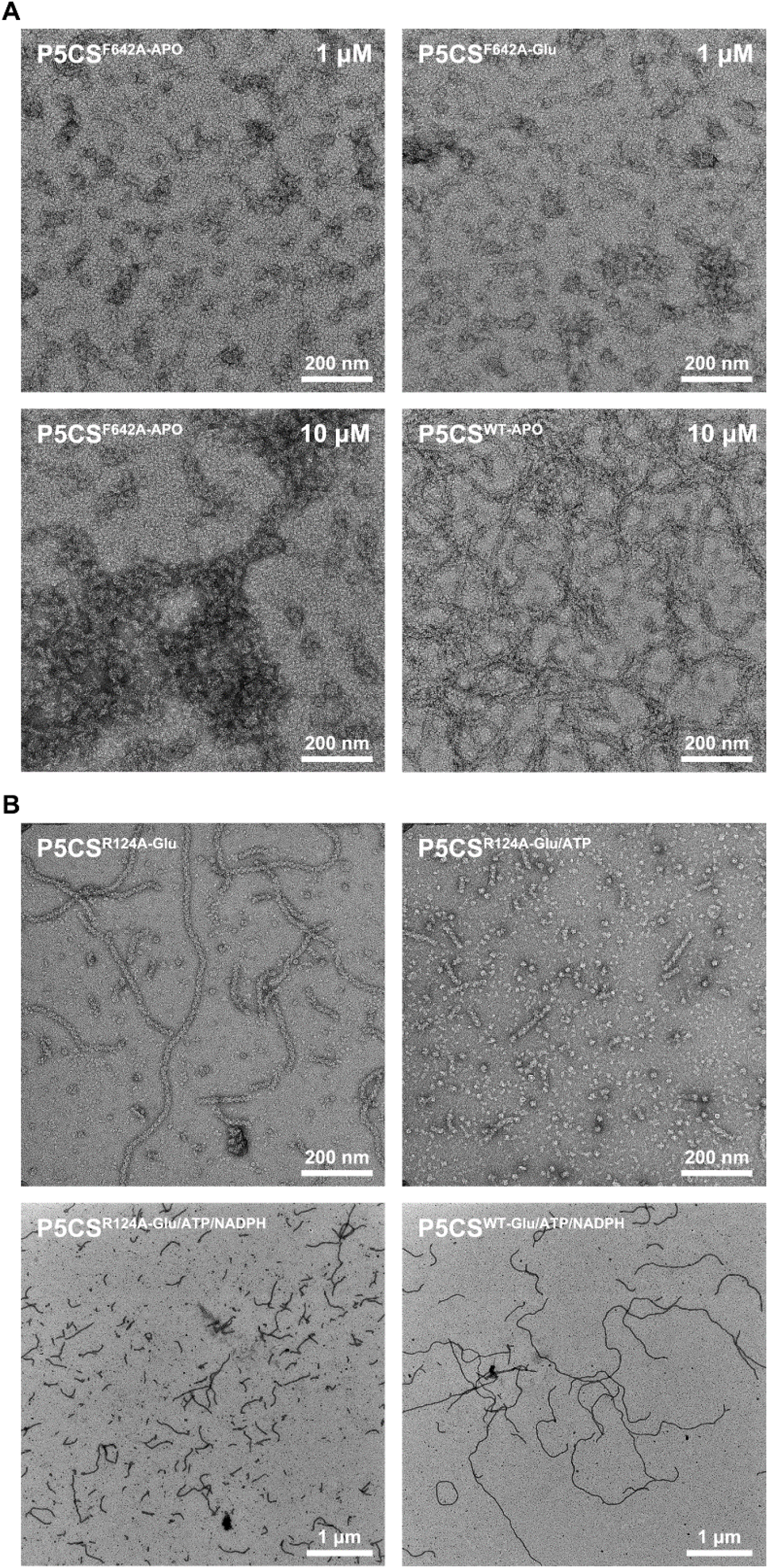
Negative staining of mutated P5CS. (**A**) Negative stain electron microscopy micrographs of P5CS^F642A^ mutation protein at APO state, this mutation will totally disrupt the filamentation of P5CS. (**B**) When the P5CS^R124A-Glu^ filament were additionally incubated with ATP, the depolymerization of P5CS^R124A^ filament were observed.

**Figure S5.**
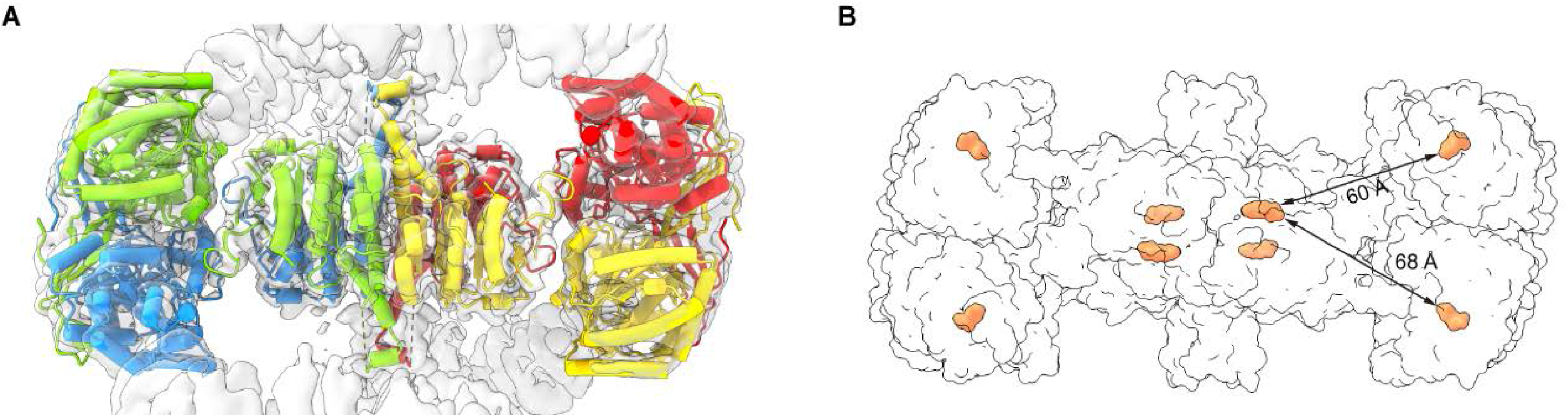
Details of P5CS filament analysis, related to Figure 5. (**A**) The tetramer model of P5CS was fitted to the cryo-EM map of filament. (**B**) Distances between the G5P (orange) in GK domain and GPR domain.

**Figure S6.**
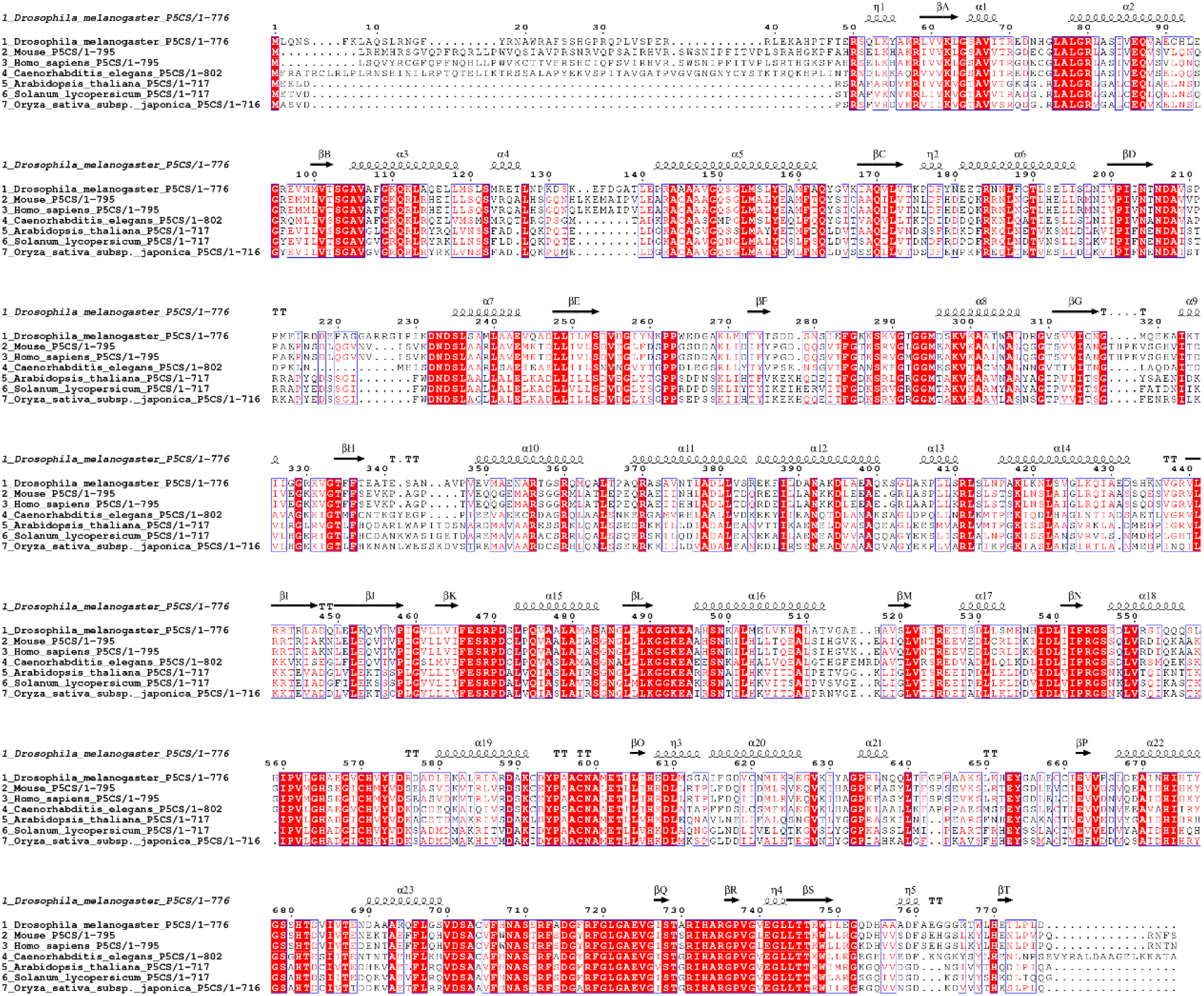
Sequence alignment of the representative P5CS enzymes. The sequence alignment of P5CS sequences of *Drosophila*, Mouse, Human, *C. elegans*, *Arabidopsis*, Tomato and Japonica rice. The conserved residues are identically shaded red and secondary structure elements are indicated above.

**Table S1.**
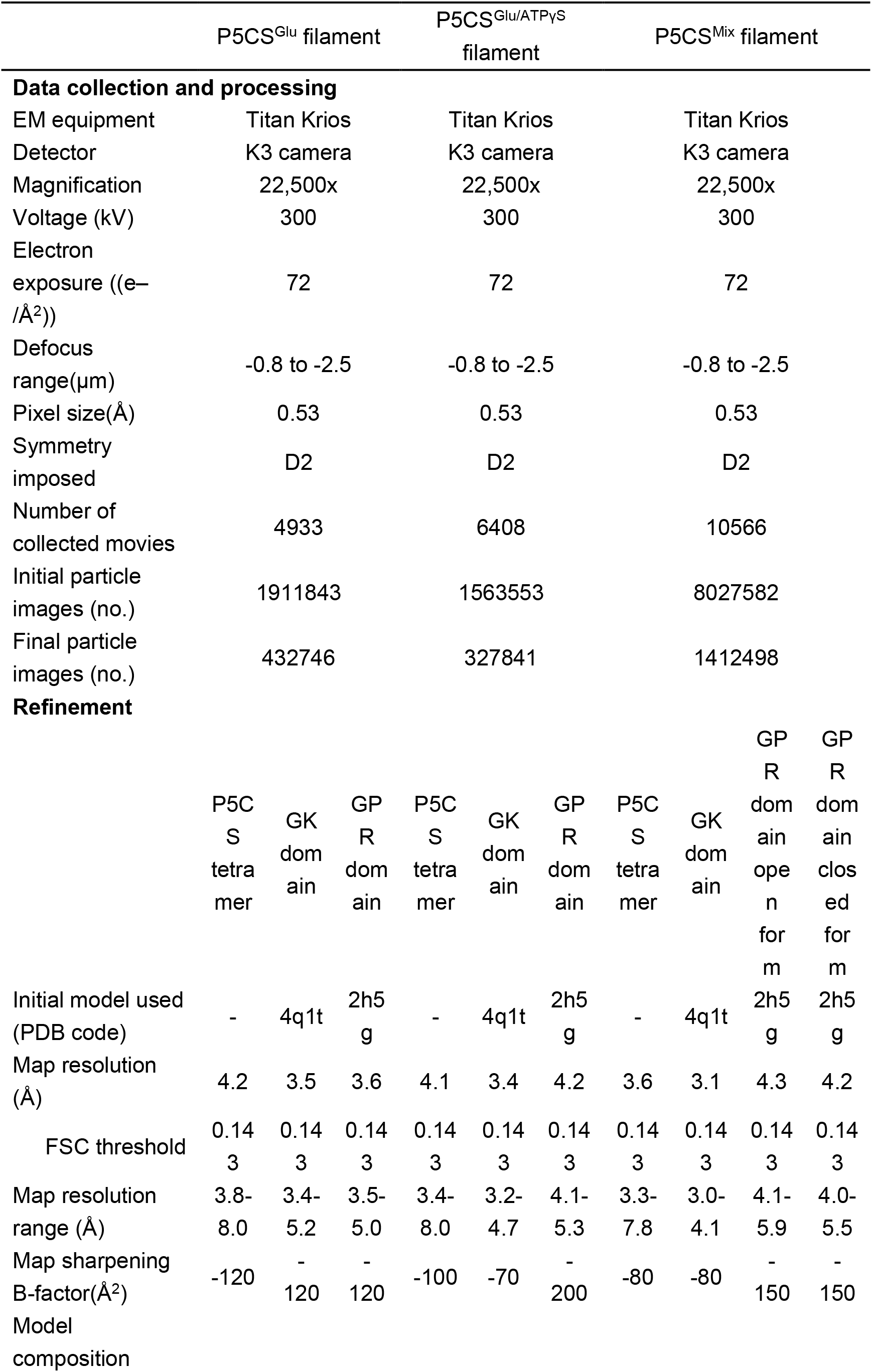
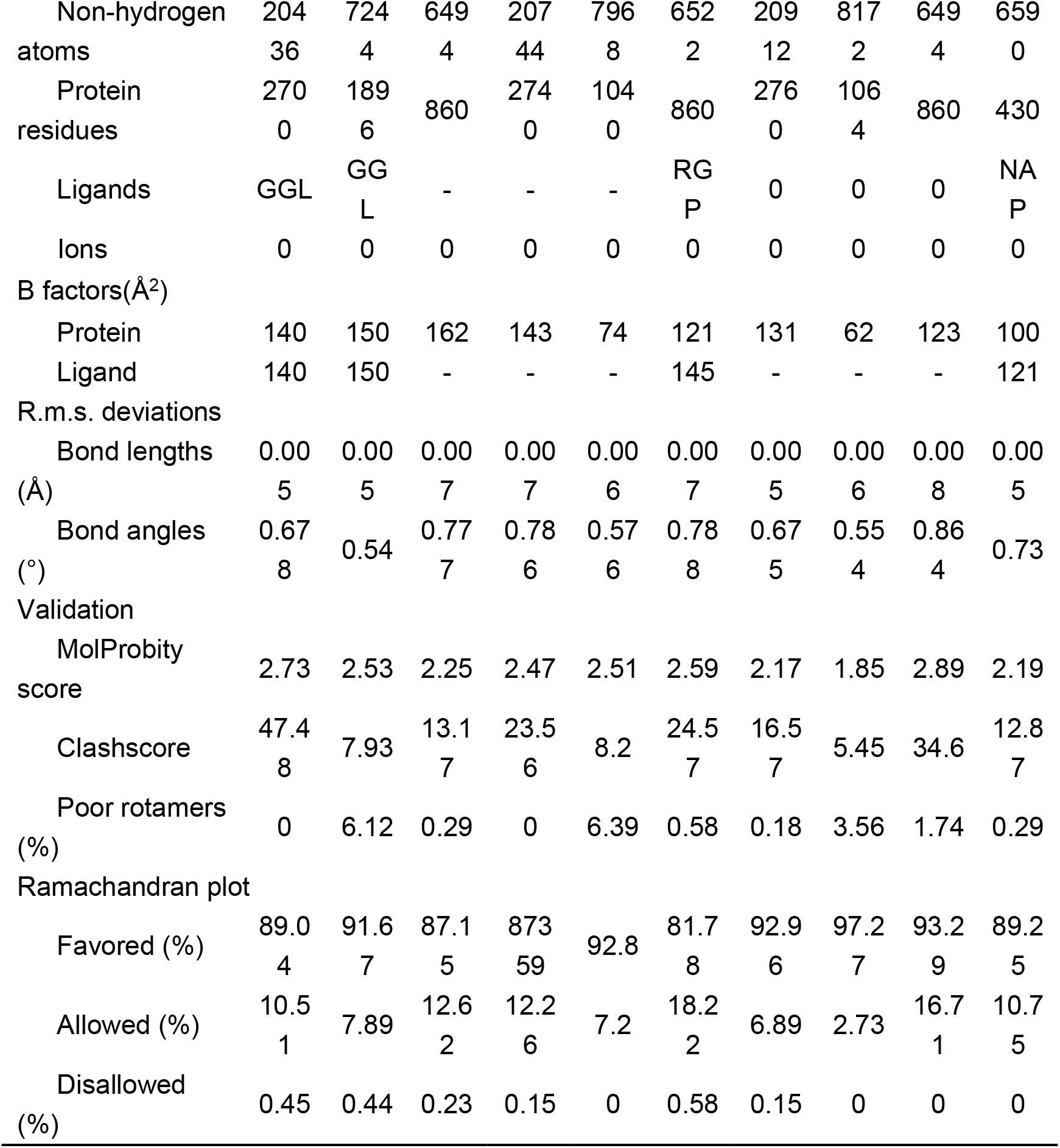
Cryo-EM data statistics.

## Legends of supplementary movie S1-S3

**Movie S1.** Morph between the consensus structures of P5CS^Glu^ filaments. Model were generated by fitting the tetramer into each classes of the 3D classification with C1 symmetry. Protomers were coloured differently. This movie implied the dynamic changes of L-Glu-bound state of P5CS filaments. Morphs between conformations were created in ChimeraX.

**Movie S2.** Morph between the consensus structures of P5CS^Mix^ filaments. Model were generated by fitting the tetramer into each classes of the 3D classification with C1 symmetry. Protomers were coloured differently. This movie implied the dynamic changes of P5CS^Mix^ filaments. Morphs between conformations were created in ChimeraX. See also Movie S1.

**Movie S3.** Morph between the consensus structures of open and closed conformations of GPR domain.

## References

1. C. A. Hu et al., Human Delta1-pyrroline-5-carboxylate synthase: function and regulation. Amino Acids 35, 665–672 (2008).

2. M. R. Baumgartner et al., Hyperammonemia with reduced ornithine, citrulline, arginine and proline: a new inborn error caused by a mutation in the gene encoding delta(1)-pyrroline-5-carboxylate synthase. Hum Mol Genet 9, 2853–2858 (2000).

3. M. R. Baumgartner et al., Delta1-pyrroline-5-carboxylate synthase deficiency: neurodegeneration, cataracts and connective tissue manifestations combined with hyperammonaemia and reduced ornithine, citrulline, arginine and proline. Eur J Pediatr 164, 31–36 (2005).

4. I. Perez-Arellano, F. Carmona-Alvarez, A. I. Martinez, J. Rodriguez-Diaz, J. Cervera, Pyrroline-5-carboxylate synthase and proline biosynthesis: from osmotolerance to rare metabolic disease. Protein Sci 19, 372–382 (2010).

5. D. L. Skidmore et al., Further Expansion of the Phenotypic Spectrum Associated With Mutations in ALDH18A1, Encoding Delta(1)-Pyrroline-5-Carboxylate Synthase (P5CS). Am J Med Genet A 155a, 1848–1856 (2011).

6. C. Marco-Marin et al., Delta(1) -Pyrroline-5-carboxylate synthetase deficiency: An emergent multifaceted urea cycle-related disorder. J Inherit Metab Dis, (2020).

7. W. Liu et al., Reprogramming of proline and glutamine metabolism contributes to the proliferative and metabolic responses regulated by oncogenic transcription factor c-MYC. Proc Natl Acad Sci U S A 109, 8983–8988 (2012).

8. Y. F. Guo et al., Inhibition of the ALDH18A1-MYCN positive feedback loop attenuates MYCN-amplified neuroblastoma growth. Sci Transl Med 12, (2020).

9. J. L. Liu, Intracellular compartmentation of CTP synthase in Drosophila. J Genet Genomics 37, 281–296 (2010).

10. M. Hunkeler et al., Structural basis for regulation of human acetyl-CoA carboxylase. Nature 558, 470–474 (2018).

11. M. C. Johnson, J. M. Kollman, Cryo-EM structures demonstrate human IMPDH2 filament assembly tunes allosteric regulation. Elife 9, (2020).

12. P. R. Stoddard et al., Polymerization in the actin ATPase clan regulates hexokinase activity in yeast. Science 367, 1039–1042 (2020).

13. C. K. Park, N. C. Horton, Structures, functions, and mechanisms of filament forming enzymes: a renaissance of enzyme filamentation. Biophys Rev 11, 927–994 (2019).

14. J. L. Liu, The Cytoophidium and Its Kind: Filamentation and Compartmentation of Metabolic Enzymes. Annu Rev Cell Dev Biol 32, 349–372 (2016).

15. X. Zhou et al., Structural basis for ligand binding modes of CTP synthase. Proc Natl Acad Sci U S A 118, (2021).

16. B. Zhang et al., The proline synthesis enzyme P5CS forms cytoophidia in Drosophila. J Genet Genomics 47, 131–143 (2020).

17. H. Gamper, V. Moses, Enzyme organization in the proline biosynthetic pathway of Escherichia coli. Biochim Biophys Acta 354, 75–87 (1974).

18. S. Cheek, K. Ginalski, H. Zhang, N. V. Grishin, A comprehensive update of the sequence and structure classification of kinases. BMC Struct Biol 5, 6 (2005).

19. Z. J. Liu et al., The first structure of an aldehyde dehydrogenase reveals novel interactions between NAD and the Rossmann fold. Nat Struct Biol 4, 317–326 (1997).

20. V. Koppaka et al., Aldehyde dehydrogenase inhibitors: a comprehensive review of the pharmacology, mechanism of action, substrate specificity, and clinical application. Pharmacol Rev 64, 520–539 (2012).

21. N. J. Zondlo, Aromatic-Proline Interactions: Electronically Tunable CH/pi Interactions. Accounts Chem Res 46, 1039–1049 (2013).

22. B. Fischer-Zirnsak et al., Recurrent De Novo Mutations Affecting Residue Arg138 of Pyrroline-5-Carboxylate Synthase Cause a Progeroid Form of Autosomal-Dominant Cutis Laxa. Am J Hum Genet 97, 483–492 (2015).

23. Z. Yang et al., Pyrroline-5-carboxylate synthase senses cellular stress and modulates metabolism by regulating mitochondrial respiration. Cell Death Differ, (2020).

